# *Setdb1*-loss induces type-I interferons and immune clearance of melanoma

**DOI:** 10.1101/2023.05.23.541922

**Authors:** Meaghan K. McGeary, William Damsky, Andrew Daniels, Eric Song, Goran Micevic, Clotilde Huet-Calderwood, Hua Jane Lou, Sateja Paradkar, Susan Kaech, David A. Calderwood, Benjamin E. Turk, Akiko Iwasaki, Marcus W. Bosenberg

## Abstract

Despite recent advances in the treatment of melanoma, many patients with metastatic disease still succumb to their disease. To identify tumor-intrinsic modulators of immunity to melanoma, we performed a whole-genome CRISPR screen in melanoma and identified multiple components of the HUSH complex, including *Setdb1*, as hits. We found that loss of *Setdb1* leads to increased immunogenicity and complete tumor clearance in a CD8+ T-cell dependent manner. Mechanistically, loss of *Setdb1* causes de-repression of endogenous retroviruses (ERVs) in melanoma cells and triggers tumor-cell intrinsic type-I interferon signaling, upregulation of MHC-I expression, and increased CD8+ T-cell infiltration. Furthermore, spontaneous immune clearance observed in *Setdb1*^*-/-*^ tumors results in subsequent protection from other ERV-expressing tumor lines, supporting the functional anti-tumor role of ERV-specific CD8+ T-cells found in the *Setdb1*^*-/-*^ microenvironment. Blocking the type-I interferon receptor in mice grafted with *Setdb1*^*-/-*^ tumors decreases immunogenicity by decreasing MHC-I expression, leading to decreased T-cell infiltration and increased melanoma growth comparable to Setdb1^*wt*^ tumors. Together, these results indicate a critical role for *Setdb1* and type-I interferons in generating an inflamed tumor microenvironment, and potentiating tumor-cell intrinsic immunogenicity in melanoma. This study further emphasizes regulators of ERV expression and type-I interferon expression as potential therapeutic targets for augmenting anti-cancer immune responses.

## Introduction

Melanoma is the deadliest form of skin cancer, and its incidence has been steadily rising(Siegel et al., 2022). Advances in immunotherapies over the last decade have led to improvements in survival of patients with advanced melanoma. However, many patients either do not respond or develop resistance to these therapies, and mortality remains high. While several mechanisms of resistance to immunotherapies have been identified through clinical and translational studies, there remains a lack of understanding as to the biology underlying a successful immune response to melanoma.

Several CRISPR screens(Griffin et al., 2021; Ishizuka et al., 2019; Manguso et al., 2017; Pan et al., 2018; Patel et al., 2017; Shalem et al., 2014) have been performed to identify and understand factors and pathways responsible for modulating the antitumor immune response, using a variety of models and with varying ranges of target pools. Here, we present a whole-genome CRISPR screen performed in a fully immunocompetent system using the syngeneic YR-G tumor model(Meeth et al., 2016), which is a UV-mutagenized murine melanoma cell line driven by common melanoma driver genes *Braf*^*V600E*^, *Cdkn2a*^-/-^ and *Pten*^-/-^, along with *Mc1r*^-/-^. Consistent with the published YUMMER1.7 cell line(Wang et al., 2017), when YR-G cells are engrafted subcutaneously into syngeneic C57BL/6J mice, tumors grow rapidly, are robustly infiltrated with immune cells, and are responsive to treatment with immunotherapies. Therefore, this model enables studies of the immune system to melanoma tumors with drivers, mutational burdens, and immune responses akin to human melanoma. Consistent with the findings of other screens, our screen identified several tumor-intrinsic pathways as critical to the anti-tumor immune response, including the Interferon-gamma pathway and MHC-I antigen processing and presentation pathway (Dubrot et al., 2022; Manguso *et al*., 2017; Patel *et al*., 2017), validating our approach.

This screen identified all members of the HUSH (HUman Silencing Hub) Complex(Tchasovnikarova et al., 2015) and its co-factors, *Setdb1(Schultz et al*., *2002), Atf7ip(Timms et al*., *2016)* and *Morc2a*(Tchasovnikarova et al., 2017) as tumor-intrinsic regulators of the anti-tumor immune response. The HUSH complex(Tchasovnikarova *et al*., 2015) consists of Periphilin (PPHLN1)(Prigozhin et al., 2020), Tasor (FAM208A)(Douse et al., 2020) and M-phase phosphoprotein 8 (MPHOSPH8)(Chang et al., 2011), and is recruited to the genome through binding DNA via MORC2A (Tchasovnikarova *et al*., 2017). Together these factors recruit SETDB1, an H3K9 methyltransferase responsible for suppressing expression of transposable elements (TEs)(Fukuda et al., 2018; Fukuda and Shinkai, 2020; Kato et al., 2018; Schultz *et al*., 2002) via tri-methylation of histone lysine 9 in genomic regions surrounding TEs. Notably, *Setdb1* is amplified in human melanoma, and its increased expression is associated with disease progression (Ceol et al., 2011). While *Setdb1* has been implicated by other screens as a regulator of anti-tumor immunity in several murine models of cancer(Griffin *et al*., 2021; Hu et al., 2021; Lin et al., 2021), this is the first study to implicate all members of the HUSH complex and its regulators. These findings add credence to these studies and strengthen our understanding of the role of *Setdb1* in cancer.

Transposable elements (TEs), and in particular endogenous retroviruses (ERVs), are noted immune-activating elements in several cancer models (Chiappinelli et al., 2015; Roulois et al., 2015). One consequence of ERV expression is activation of antiviral gene programs, including increased type-I interferon signaling and antigen presentation. This can confer additional immunogenicity to tumor cells expressing ERVs (Chiappinelli *et al*., 2015). Furthermore, when translated, ERV components can themselves be processed and presented as antigens and are then capable of stimulating robust clonotypic T cell responses (Griffin *et al*., 2021; Kraus et al., 2013; Mullins and Linnebacher, 2012; Rosato et al., 2003; Saini et al., 2020). Here we demonstrate that loss of HUSH activity via *Setdb1* knockout confers anti-viral immunogenicity that results in complete YR-G tumor clearance from immunocompetent hosts. We also find that type-I interferon signaling activated via intracellular sensing of ERV elements is critical to mounting effective anti-tumor immune responses. These findings encourage development of epigenetic therapies that de-repress ERVs towards rendering tumors increasingly immunogenic. Furthermore, they uncover elements of the underlying biology to ERV expression and regulation in melanoma.

## Results

### Whole-genome CRISPR screen identifies critical pathways regulating anti-tumor immunity in melanoma

To identify targets for anti-tumor immunity in an immunogenic model of melanoma, we performed a whole-genome CRISPR/Cas9 screen in YUMMER-G (YRG), a single-cell cloned UV-mutagenized model derived from YUMM1.G1 (Meeth *et al*., 2016; Wang *et al*., 2017). YRG cells were transduced with the *Brie* (Doench et al., 2016) genome-wide mouse CRISPR knockout library in a lentiviral vector co-expressing spCas9 and sgRNA. We engrafted 2 × 10^7^ cells subcutaneously in two host mice and allowed tumors to grow for 14 days (Fig 1A). In parallel, transduced cells were grown *in vitro* for 10 population doublings. Factors that dropped out of the screening library were compared between tumors and *in vitro* cells, revealing several critical immune pathways (Fig 1B-C). Many members of the antigen processing and presentation pathway (*Calr, Pdia3, Erap1, Abcb9, Tap1, Tap2, Tapbp1, Tapbp, Tap1, B2m, H2-D1, H2-T23, H2-Q7*) and interferon-gamma receptor pathway (*Ifngr1, Ifngr2, Jak1, Jak2, Stat1, Irf1, Nmi, Gsk3a*) dropped out specifically *in vivo*, suggesting that these pathways are required for immune evasion and tumor cell survival in an immunocompetent host (Fig 1B-C; S1A-B). Additionally, the HUSH complex (*Mphosph8, Pphln1* and *Fam208a*), and associated factors *Setdb1, Atf7ip* and *Morc2a* were also drop-out hits, consistent with recent findings in other models, but including more components of the complex (Griffin *et al*., 2021; Hu *et al*., 2021; Lin *et al*., 2021). (Fig 1B-D, Table S1) Interestingly, none of these factors represent top hits when comparing *in vitro* cells to guides to guides present immediately following library transduction, indicating that these factors are particularly required for survival *in vivo*, not just at baseline (Table S2).

**Figure 1:**
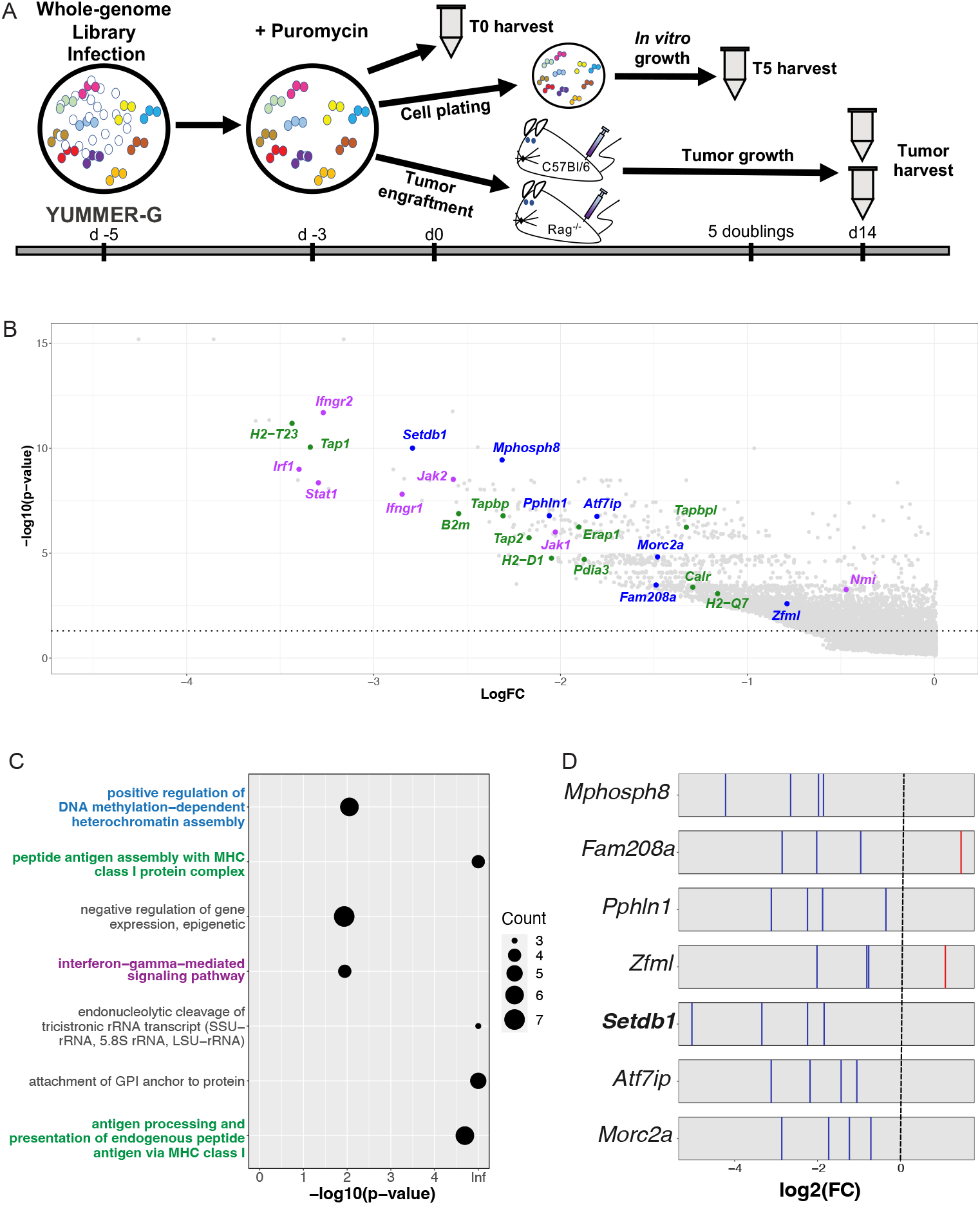
Whole-genome CRISPR screen identifies pathways critical to tumor immune evasion. 1A: Design of whole-genome CRISPR screen in YUMMER-G (YRG) cell line and tumors. 20 million cells were assayed (*in vitro*) or engrafted per experiment. 1B: Volcano plot indicating drop-out hits from the YRG^WT^ vs. *in vitro* screen conditions. Colored points include dropout members of the Interferon-gamma sensing pathway (purple), antigen processing & presentation pathway (green), and heterochromatin assembly (blue). X-axis represents fold change for each gene in YRG^WT^ vs. *in vitro* comparison, and y-axis represents log10 (p-value) for each gene in the same comparison. 1C: Significantly enriched pathways (p ≤ 0.05) represented by drop-out hits in tumors implanted in C57Bl/6J hosts vs. T5 *in vitro* condition. Identified pathways include HUSH complex members and effectors (represented by positive regulation of DNA methylation-dependent heterochromatin assembly), the Interferon-gamma sensing pathway and Antigen processing & presentation pathway. Circle size relates to number of genes identified per pathway. 1D: Distribution of guides targeting members of the HUSH complex and associated factors. Depleted guides shown in blue and enriched guides shown in red, for 4 guides per gene. p-values were calculated using MaGeCK algorithm to compare cells from tumors from C57Bl/6J hosts with cells cultured *in vitro* collected after 5 doublings.

### *Setdb1-*loss induces an immunogenic expression profile and activates the tumor immune microenvironment

To study the mechanism by which the HUSH complex mediates anti-tumor immunity, we used CRISPR to knockout the HUSH-associated H3K9 methyltransferase *Setdb1* in YRG (*Setdb1*^-/-^; SKO) (S2A). When grafted into WT C57BL/6J mice, SKO tumors are rejected (Fig 2A-B) in a CD8+ T-cell dependent manner (Fig 2C) by day 25, while WT YRG tumors reach endpoint by day 46 on average. Natural killer (NK) cells and CD4+ T cells are not required for tumor clearance (Fig S2D-E). We further investigated the tumor microenvironment by immunohistochemistry and found that SKO tumors have significantly increased CD8+ T cell infiltration (Fig 2D), but not other immune cell types (Fig 2D, Fig S2F-G). Together, these data suggest a more robust effective adaptive immune response potentiated by tumor-intrinsic loss of *Setdb1*.

**Figure 2:**
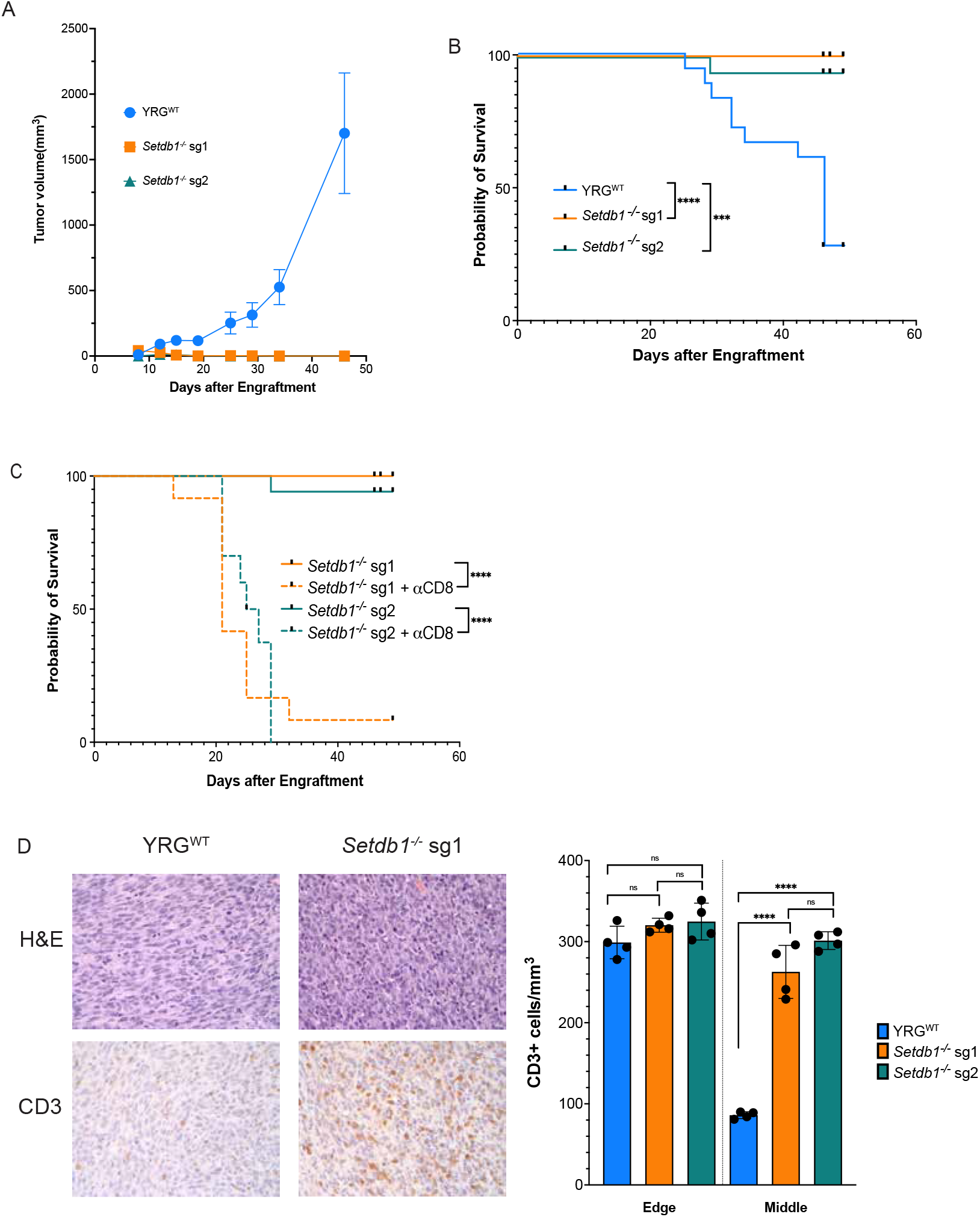
*Setdb1*-loss induces T cell infiltration and clearance of tumors in a CD8+ T-cell dependent manner. 2A: Mean tumor growth curves of YRG^WT^ tumors or *Setdb1*^*-/-*^ tumors implanted subcutaneously in syngeneic C57Bl/6J mice (1 representative cohort; N=10, 7, 8 respectively). Error bars indicate SEM. 2B: Kaplan-Meier survival curve for YRG^WT^ or *Setdb1-/-* tumors implanted in C57Bl/6J mice (2 independent cohorts; N=19, 16, 17, respectively).; Log-rank test *** p≤ 0.001; **** p≤ 0.0001. 2C: Kaplan-Meier survival curve for *Setdb1*^*-/-*^ tumors implanted in untreated or CD8-depleted host mice. N=12,11, 15, 16 per group, respectively. **** log-rank p≤ 0.0001. 2D: Representative IHC images showing T cell infiltration to YRG^WT^ and *Setdb1*^*-/-*^ tumors on d10 post-engraftment. 2E: Immune infiltration of YRG^WT^ or *Setdb1*^*-/-*^ tumors by histology, shown as % of CD3+ cells per field cells. N=4 per group; **** t-test p≤ 0.0001.

To further investigate differences between WT and SKO tumors cells we performed RNA-sequencing of WT and SKO tumor cells (Table S3). Gene set enrichment analysis (GSEA) revealed increased expression of interferon-stimulated and inflammatory response genes, including genes in the Interferon Alpha and Gamma Response HALLMARK gene sets(Subramanian et al., 2005), further indicating an inflammatory phenotype (Fig S3A). Flow cytometry analysis of WT and SKO tumor cells consistently indicated increased expression of MHC-I and antigen presentation pathway components (Fig 3A, 3D). Together these findings indicate that cells lacking *Setdb1* bear gene expression profiles consistent with tumor-intrinsic immune activation, which contributes to an inflammatory tumor microenvironment.

**Figure 3:**
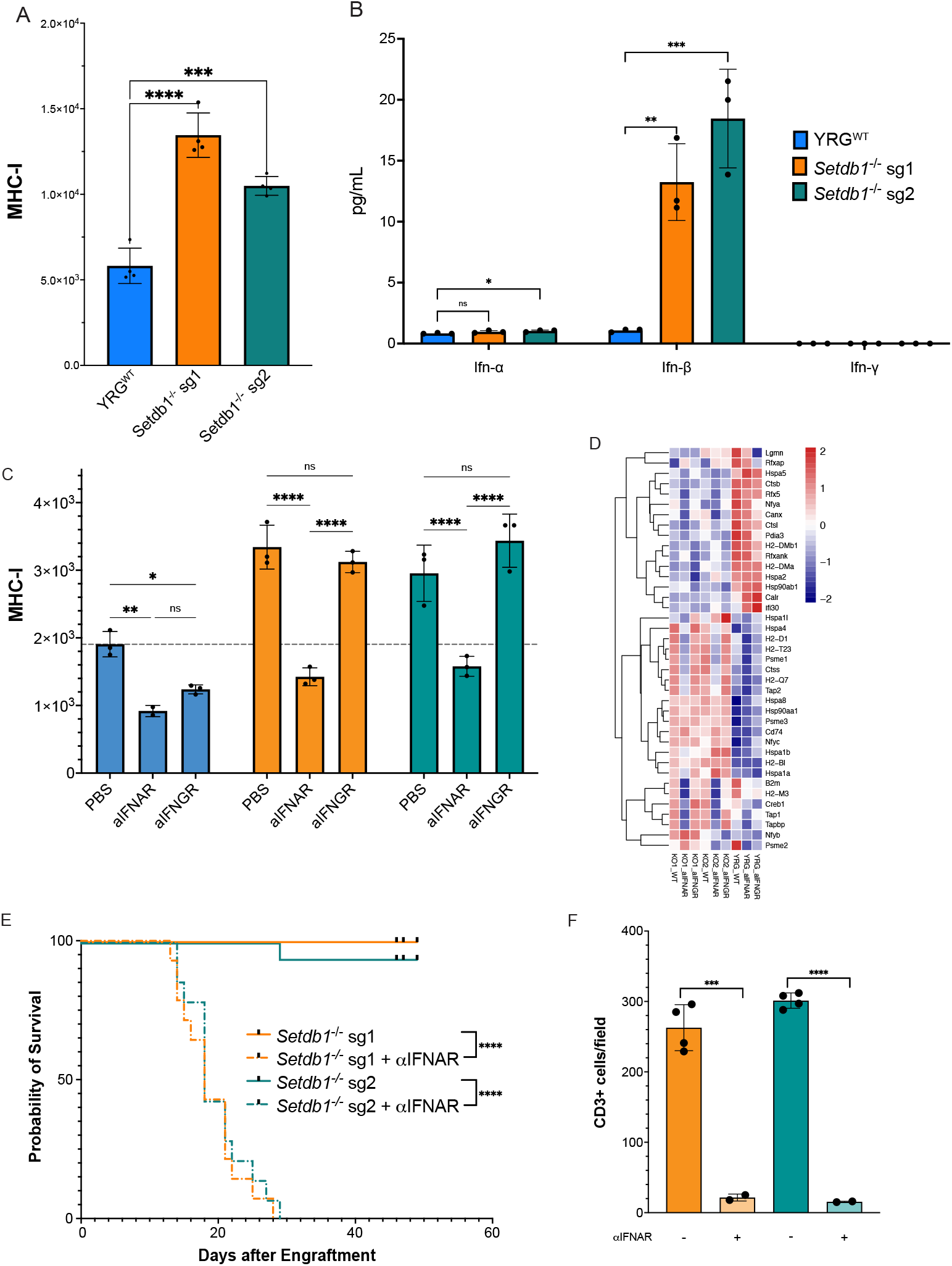
*Setdb1*^-/-^ tumor immunogenicity and clearance is dependent on type-I interferon signaling. 3A: Mean fluorescence intensity (MFI) of MHC-I expression levels by flow cytometry in YRG^WT^ or *Setdb1*^*-/-*^ cells. N=4 per group; *** indicates t-test p≤ 0.001; ****p≤ 0.0001 3B: Interferon concentrations in conditioned media of YRG^WT^ or *Setdb1*^-/-^ cells. N=3 per group; significance assessed by t-tests, *p≤ 0.05; **p≤ 0.01; ***p≤ 0.001. 3C: Mean fluorescence intensity (MFI) of MHC-I expression by flow cytometry in cultured YRG^WT^ and *Setdb1-/-* cells treated with PBS, IFNAR-blockade, or IFNGR-blockade antibodies. N=3 per group; significance assessed by t-tests, *p≤ 0.05; **p≤ 0.01; ***p≤ 0.0001. 3D: Heatmap indicating changes in MHC-I pathway gene expression between YRG^WT^ and *Setdb1*^-/-^ tumors, treated with PBS, IFNAR-blockade, or IFNGR-blockade. Colors are indicative of z-scores of normalized gene expression values between samples. 3E: Kaplan-Meier survival curves of *Setdb1*^*-/-*^ tumors untreated or treated with IFNAR blockade antibodies for duration of tumor growth. Representative of 2 experiments; N=9,9,8 per group respectively. **** indicates log-rank p≤ 0.0001. 3F: Immune infiltration of *Setdb1*^*-/-*^ tumors untreated or treated with IFNAR-blockade by histology, shown as % of CD3+ cells per field cells. N=4 per group; *** t-test p≤ 0.0005; **** t-test p≤ 0.0001.

### A tumor-intrinsic type I interferon loop is responsible for SKO cell immunogenicity

To investigate the inflammatory phenotypes we observed, we profiled cytokine secretion in SKO and YRG cells. We found that SKO cells secrete 15-20-fold more IFN-β than YRG cells, but not IFN-α or IFN-γ (Fig 3B, S3D). We hypothesized that IFN-β might be responsible for increased MHC-I expression and an inflammatory phenotype through a tumor cell-intrinsic signaling loop. To test this, we blocked the Interferon-alpha/beta receptor (IFNAR) on cells in culture with IFNAR-blocking antibody or PBS. We found that IFNAR blockade, but not IFNGR blockade, abrogates the MHC-I increase (Fig 3C) and expression of antigen-presentation and processing factors (Fig 3D), ISGs and JAK-STAT signaling in RNA-seq in SKO cells (Fig S3B-D), suggesting that loss of *Setdb1* activates a type-I interferon signaling loop in tumor cells, generating tumor-intrinsic inflammation.

IFNAR-blockade *in vivo* triggered rapid outgrowth of SKO tumors (Fig 3F), indicating that type-I interferon signaling is required for the anti-tumor immune response activated by *Setdb1*-loss. Furthermore, histology studies of tumors from mice treated with IFNAR-blockade had significantly fewer intratumoral T-cells suggesting that this type-I interferon axis is critical to recruitment of T -cells to the SKO tumor microenvironment (Fig 3F, S3E). We further profiled these tumors by single-cell RNA sequencing (scRNAseq), observing consistent microenvironment changes in tumors treated with IFNAR-blocking antibodies. Of note, SKO tumors have increased proportion of T-cells bearing an Interferon-activated gene signature (*CD8_3)*, which was decreased by IFNAR blockade (Fig S3F-G).

### SKO cells bear re-expressed ERVs which are antigenic but not immunodominant

We hypothesized that loss of *Setdb1* lead to de-repression of TEs in SKO cells, given known its roles in regulating TE expression. Analysis of our RNA-Seq data revealed increased expression of TEs, and particularly of Long Terminal Repeats (LTRs) (Fig 4A; S4A) in SKO cells. We assayed SKO and WT cells by flow cytometry for their expression of *envelope* protein, a component of MuLV-derived elements and marker for ERV expression. We found *env* expression to be strikingly increased in SKO cells (Fig 4B-C). Based on these findings, we hypothesized that *Setdb1* loss caused tumor clearance through three possible mechanisms: 1) interferon-induced inflammation of CD8+ T cells, supporting their role in anti-tumor activation; 2) interferon-induced inflammation renders tumor cells more susceptible to attacking T-cells; 3) induction of ERV antigens that T cells can recognize above those available in YRG tumors.

**Figure 4:**
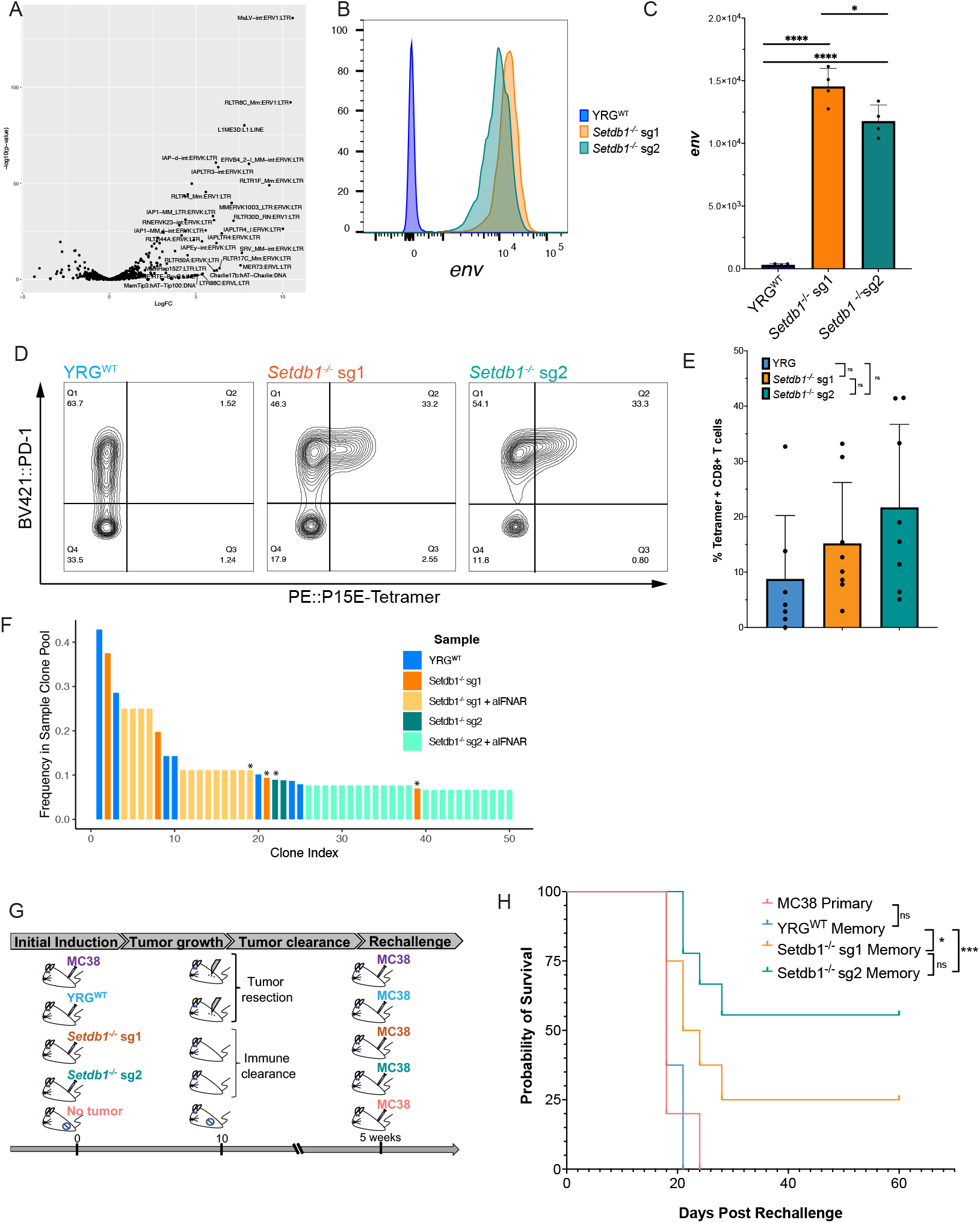
*Setdb1-*loss induces expression of ERVs, which provide substrate antigens for immune attack. 4A: Volcano plot showing upregulated transposable elements in *Setdb1*^*-/-*^ cells relative to YRG^WT^ cells. Elements shown in text have LogFC > 4 or -log10 (pval) > 20. 4B: Flow plot showing *env* expression in representative YRG^WT^ or *Setdb1*^*-/-*^ cells. 4C: MFI of *envelope* protein expression by flow cytometry in YRG^WT^ and *Setdb1*^*-/-*^ cells grown *in culture*. N=4 per group; t-tests; * p≤ 0.05; ****p≤ 0.0001 4D: Flow plots showing PD-1 expression and P15E-tetramer binding in CD8+ T cells isolated from YRG^WT^ or *Setdb1*^*-/-*^ tumors at day 12; representative samples, N=8 per group. 4E: Quantification of % tetramer positive of CD8+ T cells from YRG^WT^ or *Setdb1*^-/-^ tumors. 2 independent experiments; N=8 per group. 4F: Top 50 T cell clones identified in scRNA-seq of YRG, SKO +/-IFNAR blockade tumors, colored by sample. Data represents 3 tumors per sample. * indicates clone has matching CDR3 sequence to P15E-reactive TCR sequences from Griffin et al. 2021. 4G: Schematic of MC38 rechallenge experiment. Mice were immunized with either YRG^WT^, MC38 or Setdb1^-/-^ YRG tumors which were then either surgically resected or spontaneously cleared. 6 weeks after initial injection, mice were re-challenged with MC38 tumors and evaluated for tumor growth. N=4, 8, 4, 8, 5 per group, respectively. 4H: Kaplan-Meier survival curves of mice bearing MC38 tumors after immunization with MC38, YRG^WT^ or *Setdb1*^*-/-*^ YRG cells, which were either rejected (*Setdb1*^-/-^) or surgically resected (MC38 and YRG^WT^).

To evaluate the role of ERV antigens in rendering SKO tumors immunogenic, we used P15E-tetramers to detect env-specific CD8+ T cells specific for this *env*-derived epitope (KSPWFTTL) in the tumor microenvironment of SKO tumors (Fig 4D-E). The presence of P15E-positive CD8 T cells raises the possibility that immune clearance of SKO tumors is supported by the presence of ERV antigens. Interestingly, there is substantial variability of proportion of P15E-tetramer positive T cells present in the tumor microenvironment and these most often form a minority of infiltrating CD8+ T cells, in contrast to findings in other models (Griffin *et al*., 2021; Ye et al., 2020). Additionally, we found 25 T cells with CDR3 sequences in our scRNA-seq dataset matching published P15E-responsive CDR3 sequences (Griffin *et al*., 2021). Anticipating that exact CDR3 matches may limit detection P15E-responsive T cells, we used GLIPH (v2.0; cite) to further probe those cells whose TCRs are putatively responsive to the same epitope (see Methods), expanding the group to 78 T cells (Fig S5B). Likely due to the under-estimation of P15E-reactive T cells, we observed no trend in prevalence across samples (Fig S5C). However, these P15E-reactive clones were among the most frequent clones in the dataset, supporting a role for P15E-reactive T cells in clearance of SKO tumors (Fig 4F).

We leveraged the published finding that P15E is the immunodominant epitope in MC38 (Ye *et al*., 2020) to evaluate the functional capacity of P15E-reactive T cells to respond to SKO tumors. We grafted YRG, MC38 and SKO cells into C57BL/6J mice, and surgically removed tumors (YRG, MC38) or allowed them to clear (SKO), then grafted MC38 tumors into all mice to observe whether inducing immunological memory to P15E was sufficient to clear MC38 tumors (Fig 4G). Indeed, we found that 25-55% of mice immunized with SKO tumors could completely clear MC38 tumors upon challenge (Fig 4H), suggesting that P15E antigen-specific T cells generated during the immune response to SKO tumors provide immunity against the P15E immunodominant MC38 cell line. However, the heterogeneity of this response suggests that additional antigens, whether ERV-derived or otherwise, likely potentiate the T cell response to SKO tumors, which are not shared with MC38 tumors. To identify potential overlapping antigens, we analyzed ERV expression in publicly available MC38 RNA-seq data (Raghavan et al., 2021) and compared it with our expression data. Interestingly, we found that MC38 shares substantially more overexpressed ERV elements with *Setdb1*^-/-^ sg2 YRG cells than *Setdb1*^-/-^ sg1 YRG cells, consistent with finding that *Setdb1*^-/-^ sg2 YRG tumors provoke a better recall response to MC38 tumors upon re-challenge (Fig S4B-C). we also compared expression levels of cancer-testis antigens (Almeida et al., 2009) in YRG, SKO and MC38 cells and found no overlap (Fig S4E), further supporting the ERV antigen hypothesis.

### Discussion

Here we present the results of a whole-genome CRISPR/Cas9 screen performed in a fully immunocompetent melanoma mouse model, which has uncovered a role for several critical pathways. Our findings converge on pathways identified by other melanoma CRISPR screens (Dubrot *et al*., 2022; Griffin *et al*., 2021; Ishizuka *et al*., 2019; Manguso *et al*., 2017; Pan *et al*., 2018; Patel *et al*., 2017). It is clear from our work, and that of others, that antigen presentation, supported by IFN-γ sensing, is a primary modulator of resistance to T cell responses, and potentially of susceptibility to NK cell attack (Manguso *et al*., 2017; Pan *et al*., 2018; Patel *et al*., 2017) (Fig. 1B-C). We find that loss of factors from both these pathways increased susceptibility of tumor cells to death in immune-competent hosts. Loss of antigen presentation machinery has been identified as a mechanism of resistance in a variety of melanoma models and in patient populations, allowing tumor cells to evade T cell attack. a role for NK cells in combatting this resistance by NK cell-mediated detection of tumor cells in response to their lack of surface MHC-I has also been proposed (Moretta et al., 2001). We hypothesize that our findings are indicative of this phenomenon in modulating the fitness landscape of YRG tumors. Furthermore, when taken with the findings of (Benci et al., 2019; Benci et al., 2016) it is possible that loss of interferon-gamma sensing contributes to the recruitment and activation of innate immune cells such as NKs and Innate lymphoid cells (ILCs) that affect efficient and antigen-independent tumor killing (Benci *et al*., 2019). Adaptive loss of alternative MHC-I molecules such as H2-Q7 and H2-T23 are consistent with the findings of (Manguso *et al*., 2017), and further support this hypothesis. Taken together these results validate our CRISPR screening platform and findings, which newly identified a role for the entire HUSH complex in anti-tumor immunity to melanoma.

Tumor loss of the HUSH complex-associated methyltransferase *Setdb1* potentiated a striking anti-tumor T cell response independent of NK cells, indicative of multiple types of immune killing occurring within heterogeneous tumors. Accordingly, here we describe the induction of novel ERV antigens and a type-I interferon signaling loop that results from *Setdb1* loss in tumor cells. We further find that type-I interferon secretion is required for effective T cell trafficking to tumors and efficient and durable clearance of *Setdb1* tumors by CD8+ T cells. Finally, we provide evidence that some SKO cell-reactive T cells are specific to ERV antigens induced through *Setdb1* loss and likely have anti-tumor efficacy in other tumor types, including MC38. This study strongly supports the anti-tumor impact of type-I interferon signaling, particularly in the context of CD8+ T cell killing. Specifically, IFN-β is required for upregulation of MHC-I in *Setdb1*^*-/-*^ tumors, increasing tumor cell immunogenicity. Furthermore, we found that blocking type-I interferon signaling in host mice prevented clearance of *Setdb1*^-/-^ tumors, indicating that this signaling loop not only augments anti-tumor immunity but is in fact central to it. Several studies have shown similar findings, in that epigenetic upregulation of ERVs triggers an anti-viral response including type-I interferons and augmenting anti-tumor immunity (Chiappinelli et al., 2016; Roulois *et al*., 2015).

A previous study evaluating the impacts of *Setdb1*^*-/-*^ loss in B16 melanoma found no upregulation of a type I interferon response and hypothesized that anti-tumor immunity was mediated entirely through expression of novel antigens (Griffin *et al*., 2021). One possible explanation for this is upregulation of different viral elements in the B16 model (relative to YRG), resulting in lack of innate sensor activation. Griffin et al. found that *Trim28* knockout, but not *Mpp8* knockout conferred a similar survival benefit to *Setdb1* knockout in B16 cells, suggesting that *Trim28*-*Setdb1* co-mediated H3K9 trimethylation might be the primary mechanism suppressing ERVs in B16 melanoma.

Interestingly, they found a role for the HUSH complex over *Trim28* activity in a lung cancer model (LLC)(Griffin *et al*., 2021). Our CRISPR screen findings strongly implicate the activity of the HUSH complex as important to suppressing ERVS to prevent activation of anti-tumor inflammation in YRG (Fig. 1C-D). It is also possible that B16, LLC and YRG melanomas have different ERVs de-repressed on *Setdb1* loss, resulting in differential innate sensing behaviors and interferon induction. Alternatively, there may be additional tissue or cell-specific mechanisms that cooperate with these H3K9me mechanisms, driving their differential patterns.

We further show that YRG melanomas lacking *Setdb1* are efficiently and durably cleared by cytotoxic T cells, leading to cure (Fig. 2A-C). This is in striking contrast to the findings of Griffin et al. in B16 melanoma, *Setdb1*-loss coupled with immune checkpoint blockade treatment is sufficient to extend survival, but not to induce tumor cure (Griffin *et al*., 2021). While these differences could be mediated by differences in relative tumor dosages, we found that *Setdb1*^*-/-*^ tumor engraftments of 1*10^7^ cells are durably cleared, suggesting this might not be the case (data not shown). An alternative hypothesis is the relative antigen load of B16 and YRG tumors; YRG tumors bear a substantial UV-induced mutation burden(Alexandrov et al., 2020) (data not shown), which likely leads to an increased antigen load and may explain its increased immunogenicity compared with B16 tumors. Indeed, we found that B16 tumors contain a proportion of P15E-tetramer positive T cells resembling that of SKO tumors, suggesting that additional components (such as interferon signaling or additional antigens) may be necessary for complete tumor clearance (Fig. S4E). We hypothesize that the breadth of available antigens in SKO tumors might account for the observed difference in immune response, which might be evaluated in the future using UV-irradiated cell lines bearing *Setdb1*-loss (Wolf et al., 2019). We found that 25-55% of SKO tumor-experienced mice were competent to clear MC38 tumors suggesting a greater overlap of antigens to SKO tumors (Fig. 4E-F). The correlation between ERV expression level in SKO lines, subsequent tetramer positivity intratumoral CD8+ T cells, and efficacy of memory to MC38 is interesting to note, and further supports that ERV-detecting T cells are the primary effector in this context. Ye et al. found that P15E is an immunodominant antigen in MC38, but not sufficient to mediate tumor clearance(Ye *et al*., 2020). Comparison of expressed ERVs revealed several candidate factors that may underly the cross reactivity of MC38 and SKO immune response (Fig S4B). Importantly, MC38 and SKO cells do not share potential neoepitopes or over-expression of any known cancer-testis antigens, further supporting a role for ERV-derived antigens. (Fig S4E). Functional evaluation of these antigens, coupled with investigation of type I interferon responses will undoubtedly uncover the relative roles of these phenotypes in mediating responses competent to result in tumor clearance.

Together, these findings further support therapeutic targeting of *Setdb1* to reactivate ERV expression, which we show is both sufficient to contribute novel antigens and type-I interferon signaling to the immune landscape. These findings agree with a recent study that showed ERVs could be re-activated through loss of the H3K4 histone demethylase *Kdm5b*, which in turn requires *Setdb1* for ERV suppression. Loss of *Kdm5b* similarly activates type-I interferon signaling through sensing of cytosolic DNA via *Sting*, suggesting this may also underly innate signaling in our *Setdb1*^*-/-*^ cells (Zhang et al., 2021). Study of the co-factors of *Setdb1*, including *Kdm5b*, the HUSH complex, *Trim28, Atf7ip*, and *Morc2a* are likely to further elucidate the mechanisms by which ERVS are repressed and can be therapeutically re-activated in melanoma and other cancers. In addition to implicating *Setdb1* as a therapeutic target, this work supports targeting ERV expression as a strategy for augmenting the anti-tumor immune response and improving patient responses to immunotherapy (Chiappinelli *et al*., 2015; Griffin *et al*., 2021; Gu et al., 2021; Kraus *et al*., 2013; Krishnamurthy et al., 2015; Mullins and Linnebacher, 2012; Ye *et al*., 2020; Zhou et al., 2015). Regulation of ERV expression in human cells is poorly understood but will likely be critical for developing therapies that induce anti-tumor immunity through increase of inflammatory signaling and provision of additional potent antigens to the tumor microenvironmental milieu.

## Data Availability

RNA-sequencing data and single-cell RNA-sequencing data will be made available on Gene Expression Omnibus upon publication.

## Acknowledgements

We gratefully acknowledge members of the Bosenberg Lab and Yale Cancer Center research community for their helpful comments and suggestions. This work was supported by NCI F31CA243212 and F99CA253767 (to MKM), as well as P50CA121974, U01CA238728, U01CA233096 (to MWB) and a Melanoma Research Alliance Team Science Award (to MWB and WD). GM is supported by a NIAID-funded fellowship T32AR007016-47 to Yale Department of Dermatology. GM has been supported by the Dermatology Foundation and American Skin Association. We also thank the Yale Stem Cell Center Genomics Core, Yale Center for Genome Analysis, Yale Flow Cytometry Core and Yale Pathology Tissue Services (YPTS), and Yale Cancer Center Functional Genomics Core for their expert assistance and services.

## Author Contributions

MKM, WD, AD, GM, SP, DAC, SK, BET, AI, and MWB contributed to the design of this study. Experiments were performed by MKM, WD, AD, ES, GM, CHC, HJL, and SP. MKM, WD, AD, CHC, SP, DAC, BET, AI, and MWB analyzed and interpreted data. MKM, WD, AD, GM, DAC, BET, and MWB contributed to writing and figure design. MWB and WD supervised the study.

## Methods

### Cell Lines

Cell lines were grown in Complete Media (DMEM/F12 (Gibco), 1X NEAA (Gibco), 1X PenStrep (Gibco), 10% FBS (Gibco) at 37°C in 5% CO_2_, maintained at low passages and passaged using 0.25X Trypsin EDTA (Invitrogen). Cells lacking *Setdb1* were achieved by CRISPR/Cas9 gene editing using sgRNAs selected from the Brie library(Joung et al., 2017). The sgRNAs were introduced into the px458 vector(Ran et al., 2013), which was verified by Sanger sequencing and transiently transfected into target cell lines via Lipofectamine 3000 (Invitrogen) or TransIT-LT1 (Mirus) reagents according to manufacturer instructions. 48 hours after transfection, cells were single-cell sorted using BD-FACS Aria on GFP positivity and grown out. Knockouts were validated by western blotting using GAPDH and compared with control, un-transfected cells (Fig. S2A). Two cell clones (sg1-G2 and sg2-G4) were selected for further studies and are labelled throughout sg1 or sg2, respectively.

### Mouse Experiments

2 × 10^6^ cells were injected subcutaneously into the right hindlimb of 9 to 12-week-old male C57BL/6J mice (Jackson Laboratories). Tumors were measured using a caliper twice weekly following appearance of tumors, and tumor volumes were calculated using the equation 0.5233* (length)* (width)* (depth). Mice were sacrificed when tumors reached an endpoint volume of 1cm^3^ or ulcerated, according to Yale IACUC-approved animal protocols. For depletion experiments, mice were treated twice weekly with intraperitoneal injection of 200ug aCD4 (GK1.5), aCD8 (2.43), or aNK1.1 (PK136) antibody (BioXcell), beginning at least 3 days prior to tumor engraftment and for the duration of tumor outgrowth or until no palpable tumor could be observed. For blockade experiments, mice were treated twice weekly with intraperitoneal injection of 200ug aIFNAR (MAR1-5A43; BioXcell) beginning at tumor engraftment and continuing as described above. Rosa26^Cas9-EGFP^ (Jax #026179)(Platt et al., 2014) and C57BL/6J-TgN (pPWL88puro)2Ems (Linnell et al., 2001) (Jax #002355) mice were bred to produce host mice heterozygous for each allele. These mice express both Cas9 and Puromycin-resistance, ensuring tolerance to any potential antigens derived from these proteins in engrafted tumor cells and served as hosts for the CRISPR screen.

### Immune Memory Experiments

2 × 10^6^ YRG or SKO cells or 1 × 10^6^ MC38 cells were injected subcutaneously into the right hindlimb of 9–12-week-old male C57BL/6J mice (Jackson Laboratories). 10 days after injection, YRG and MC38 tumors were surgically removed using sterile technique according to Yale IACUC-approved surgical protocol. Mice were allowed to recover for at least 4 weeks. Once tumors had cleared (SKO) or surgical sites had healed (YRG and MC38), SKO, YRG and MC38 ‘memory’ mice or age-matched naïve mice were each re-challenged with 1 × 10^6^ MC38 cells, and tumors were monitored as described above.

### CRISPR screening

For whole-genome CRISPR screen, 2 × 10^7^ cells per experiment were treated with *Brie* lentivirus pool at a MOI of 0.3 as described(Joung *et al*., 2017). Cells were selected with puromycin for 72 hours, then 2 × 10^7^ cells were engrafted in Rosa26^Cas9-EGFP/+^ TgN (pPWL88puro)2Ems^+/-^ host mice (described above). Tumors were measured twice weekly using a caliper and removed when they reached an endpoint of 1 cm^3^ according to Yale-approved IACUC protocols. In parallel, following puromycin selection, 2 × 10^7^ cells were harvested before and after propagation for 4 doublings. Cells were also reserved after puromycin selection as a baseline control for guides present in the library. Genomic DNA was isolated from tumors or cultured cells using Qiagen Blood & Tissue kit, amplified as described (Joung *et al*., 2017), and deep sequenced using NovaSeq sequencer at the Yale Center for Genomic Analysis. Data was analyzed using MaGEcK (Li et al., 2014) (v. 0.5.6) with default parameters. Pathway analysis was performed on drop-out hits with a p-value < 0.05, using the MaGEcKFlute package (1.14.0) EnrichAnalyzer command with the HyperGeometric test and GO Biological Processes gene sets(Wang et al., 2019). Pathways with adjusted p-values <0.05 were considered significant. Plots were generated using ggplot2 (3.3.5) and ggrepel (0.9.1) packages.

### *In vitro* cell treatments

1×10^6^ cells were plated in complete media and incubate in culture for 24 hours to allow them to adhere. Cells were then treated with PBS or aIFNAR (listed above) for 72 hours (with treated media replaced every 36 hours) and analyzed using western blotting, flow cytometry, RNA sequencing or cytokine assays as described.

### Flow cytometry

Cells in culture were trypsinized (0.25X Trypsin-EDTA, Invitrogen) and pelleted, stained with antibodies to MHC-I (H2^dB^/H2^kB^; 28-8-6; Biolegend) and/or ab573(Evans et al., 2014) (1:2 hybridoma supernatant) antibodies for 20 minutes at 4°C, then stained with secondary (anti-mouse IgM) and live-dead markers (Life Technologies) for 20 minutes, washed twice with FACS buffer and analyzed on BD LSRII or BD Symphony flow cytometric analyzers. Data was analyzed using FlowJo software (v 10.8.1). Tumors were harvested from host mice 10 days after injection. Samples were dissociated in dissociation buffer (RPMI + 10% FBS + 1mg/mL collagenase + 1X DNase) rotating at 37°C for 30 minutes, then pushed through 70uM filter, treated with ACK buffer, quenched with RPMI+FBS and washed with FACS buffer (PBS + 1% FBS). Staining was performed using antibodies to CD45 (30-F11), CD4 (GK1.5/RM4-5), CD8 (53-6.7), Nkp46 (PK136), F4/80 (Bm8), CD11C (N418), CD279 (29F.1A12), MHC-I (28-8-6), and CD44 (IM7) antibodies, H2-Kb KSPWFTTL tetramer (MBL Laboratories/NIH Tetramer Core), and Fixable Live/Dead Aqua Stain (Life Tech) in PBS at 4°C for 30 minutes, incubated in fixation buffer (BioLegend) for 10 minutes at room temperature, and maintained overnight at 4°C. Samples were run next day on LSRII flow cytometric analyzers and data analyzed using FlowJo software (v. 10.8.2). Gating schemes are shown in Fig S5.

### Cytokine assays

1×10^6^ cells were plated in culture in complete media and incubated for 24 hours to allow adherence. Complete media was replaced with serum-free media for 24 hours, and then removed from cells and spun down at 13000 RPM for 5 minutes. Supernatant samples were shipped on dry ice and evaluated for cytokine concentrations by Eve Technologies.

### RNA sequencing

1×10^6^ cells were pelleted, and RNA isolated using RNeasy kit, supplemented with QIAshredder and DNase treatment (Qiagen). RNA concentration and fragment sizes were evaluated for QC using Agilent BioAnalyzer or TapeStation, then library prep performed and sequenced using Ilumina HiSeq4000 at the Yale Stem Cell Center Genomics Core. Reads were aligned using BWA (Li, 2013) (version 0.7.17) to mm10. Gene expression values were counted using HTseq (Anders et al., 2015) (version 0.13.5) and differential expression was performed by edgeR (McCarthy et al., 2012) (version 3.36.0). For analysis of transposable elements, RepEnrich was used to align to RepeatMasker (mm10) reference and generate counts for each element. Simple repeats were filtered out of results, and differential expression was performed for ERV and gene expression using edgeR (McCarthy *et al*., 2012) as above. Plots were produced using pHeatmap (v 0.1.12) and ggplot2 (3.3.5) in R. Gene set enrichment analysis (Subramanian *et al*., 2005) was performed using desktop software (v. 4.1.0) and HALLMARK gene sets downloaded from MSigDB (Liberzon et al., 2015). MC38 expression datasets were downloaded from Gene Expression Omnibus (GSM5221633)(Raghavan *et al*., 2021) and processed as described above for gene expression and ERV expression analysis. Cancer testis antigens gene names were downloaded from the CTdatabase (Almeida *et al*., 2009) and converted to mouse gene names using DAVID if possible. Venn diagrams were generated using the venn () function from gplots for R (v. 3.1.3).

### Immunohistochemistry

Tumors and matching spleens were harvested at noted time points, fixed in 10% buffered formalin and cassettes generated by the Yale Pathology Histology Tissue Resource. Staining was performed by Yale Pathology using antibodies to CD3 (clone), F4/80 (clone) and FOXP3 (clone) or Hematoxylin and Eosin. Scoring was performed by a board-certified pathologist (MWB) in a blinded fashion for number of positive cells per mm^3^.

### Single Cell RNA sequencing

s at day 10 post-injection and dissociated on ice in digest buffer (RPMI + 2% FBS, 100U/ml collagenase (Sigma) + 1X DNAse I (Sigma)). Samples were filtered through a 70uM strainers, pelleted, and treated with ACK Buffer to lyse red blood cells. Single-cell suspensions were spun, pelleted, and stained with TotalSeqC Hashtag antibodies (BioLegend) by tumor ID then pooled by sample. Samples were stained with antibodies to CD45, CD3e, CD8, Fixable Live/Dead Aqua (Life Tech), and sorted using BD FACS Aria system by the Yale Flow Cytometry Core. Cells were pooled in the following groups/ratios using cell counts from the cytometer: CD45-(25%); CD45+ CD3-(25%); CD8+ (50%). 5000 cells per sample were loaded onto the 10X Chromium Library. Preparation was performed using 10x Genomics reagents according to the manufacturer’s instructions by the Yale Center for Genome Analysis (YCGA) and passed quality control. Libraries were sequenced using an Illumina NovaSeq at the YCGA.

### Single Cell RNA sequencing Analysis

Samples were preliminarily processed using the Cellranger software suite commands cellranger mkfastq to generate fastq files, and cellranger multi to both align gene expression reads to the mm10 reference genome, assemble V (D)J contigs, generate a filtered gene expression matrix and make clonotypes calls. Cellranger aggr was then used to generate a single dataset from all samples sequenced with no normalization. Aggregated counts table was analyzed using the Seurat package for R (v. 4.3.0). Briefly, counts were read in as a Seurat object, and cells with less than 25 transcripts were filtered from the dataset. Expression values were log-normalized, variable genes detected, and values were scaled to number of UMIs. Principle component analysis was performed, and top 100 PCs were used to find nearest neighbors, perform clustering, and run dimensional reduction using the UMAP algorithm according to standard Seurat workflow. To determine which cells were most likely real, healthy cells (as opposed to dying cells or empty droplets), we identified the lowest-transcript clusters with a median number of transcripts per cell (median nFeature_RNA) <100. We performed differential expression to identify gene markers of nFeature_RNA clusters with median nFeature_RNA >100 using the FindMarkers () function. We selected markers present in >80% of non-low-nFeature_RNA clusters and <25% of low-nFeature_RNA clusters, termed ‘good.genes’, and determined the median expression value of these ‘good.genes’ for every cell in the dataset. Cells with a good.gene median ==0 were removed from the dataset. Normalization, Variable gene detection, scaling, PCA, Neighbor analysis, clustering and dimensional reduction were re-run on remaining cells. Conventional cell-type markers were used to characterize clusters, and T cell clusters were subsetted, and re-analyzed by the workflow described above. scRepertoire for R was used to analyze clonotype data. Briefly, contig annotations were combined across samples and associated with the gene expression Seurat object. HTO-Demux was used to make hashtag calls and determine which animal each T cell clonotype was derived from. The most frequently appearing clones were identified and CDR3 sequences were compared with those published by Griffin et al. as P15E-responsive. To expand this analysis, we used GLIPH (v2) to identify clusters of T cell clonotypes likely responding to the same antigen and used a custom python script to search for identified patterns in P15E-responsive clonotypes. We then expanded our list of putatively P15E-responsive T cell clonotypes to include these patterns.

## Supplemental Information

Figure S1: CRISPR screen guide distributions, related to Figure 1

A-B) Guides targeting noted Interferon-gamma and antigen processing/presentation pathways, respectively. Depleted guides shown in blue and enriched guides shown in red, for 4 guides per gene. p-values were calculated using MaGeCK algorithm to compare cells from tumors from C57Bl/6J hosts with cells cultured *in vitro* collected after 5 doublings.

Figure S2: Validation and characterization of *Setdb1*^*-/-*^ cell lines, tumors, and microenvironments, related to Figure 2.

A: Western blot indicating loss of SETDB1 expression in *Setdb1*^*-/-*^ single-cell clones. GAPDH used as a loading control.

B: Spider plots showing tumor volume in untreated host mice by tumor genotype, associated with figure 2A.

C: Spider plots showing tumor volume in host mice treated with aCD8 antibodies, over time by tumor genotype, associated with figure 2B.

D: Kaplan-Meier survival curves for *Setdb1*^*-/-*^ tumors in host mice treated with aCD4 antibodies.

E: Kaplan-Meier survival curves for *Setdb1*^*-/-*^ tumors in host mice treated with aNK1.1 antibodies.

F: Immune infiltration of SKO or YRG tumors assessed by flow cytometry as described (Methods). Representative of 3 tumors per group.

G: Immune infiltration of SKO or YRG tumors +/-IFNAR blockade by scRNA-seq. Cell groups identified through expression of known marker genes as described in Methods.

Figure S3: Evaluation of differences in SKO tumor gene expression and microenvironment profiles, related to Figure 3:

A: Gene set enrichment analysis for significantly over-expressed genes in Setdb1^-/-^ cells vs. YRG. Bars represent NES score for each pathway.

B-C: Heatmap indicating z-scores for members of the JAK/STAT pathway (B) and Ifn-a pathway (C) in YRG and *Setdb1*^*-/-*^ cells, untreated or treated with aIFNAR or aIFNGR antibodies, as noted.

D: Concentration of cytokines in supernatant media of YRG or *Setdb1*^*-/-*^ cells in culture. E: Representative histology images of T cell infiltration (CD3) SKO tumors +/-IFNAR blockade at the tumor edge and middle.

Figure S4: Characterization of possible SKO tumor antigens, related to Figure 4. A: Pie chart showing proportion of upregulated ERVs by TE class.

B: Venn-diagram indicating overlapping over-expressed ERVs in MC38 and *Setdb1*^*-/-*^ cells relative to YRG.

C: Heatmap indicating normalized gene expression counts of ERV elements significantly over-expressed in *Setdb1*^*-/-*^ cells, in SKO, YRG and MC38 cell lines.

D: Heatmap indicating normalize gene expression counts of cancer testis antigens in SKO, YRG and MC38 cell lines.

E: Representative histogram indicating % P15E tetramer-positive CD8+ T cells in *Setdb1*^*-/-*^, YRG, MC38 and B16 tumors.

Figure S5: Characterization of P15E-tetramer positive T cells in flow and bioinformatic data, related to Figure 4

A: Gating scheme for flow cytometric analysis of tetramer positivity.

B: Putative P15E T cell clones depicted on UMAP plot of T cells observed in scRNA-seq dataset as having productive TCR sequences.

C: Frequency of exact (P15E) and predicted epitope-matching (putative P15E) T cell clones by sample in scRNA-seq data.

