## Supplementary figures and images for "*Setdb1*-loss induces type-I interferons and immune clearance of melanoma"

### Supplemental Figures

#### Supplemental Figure 1

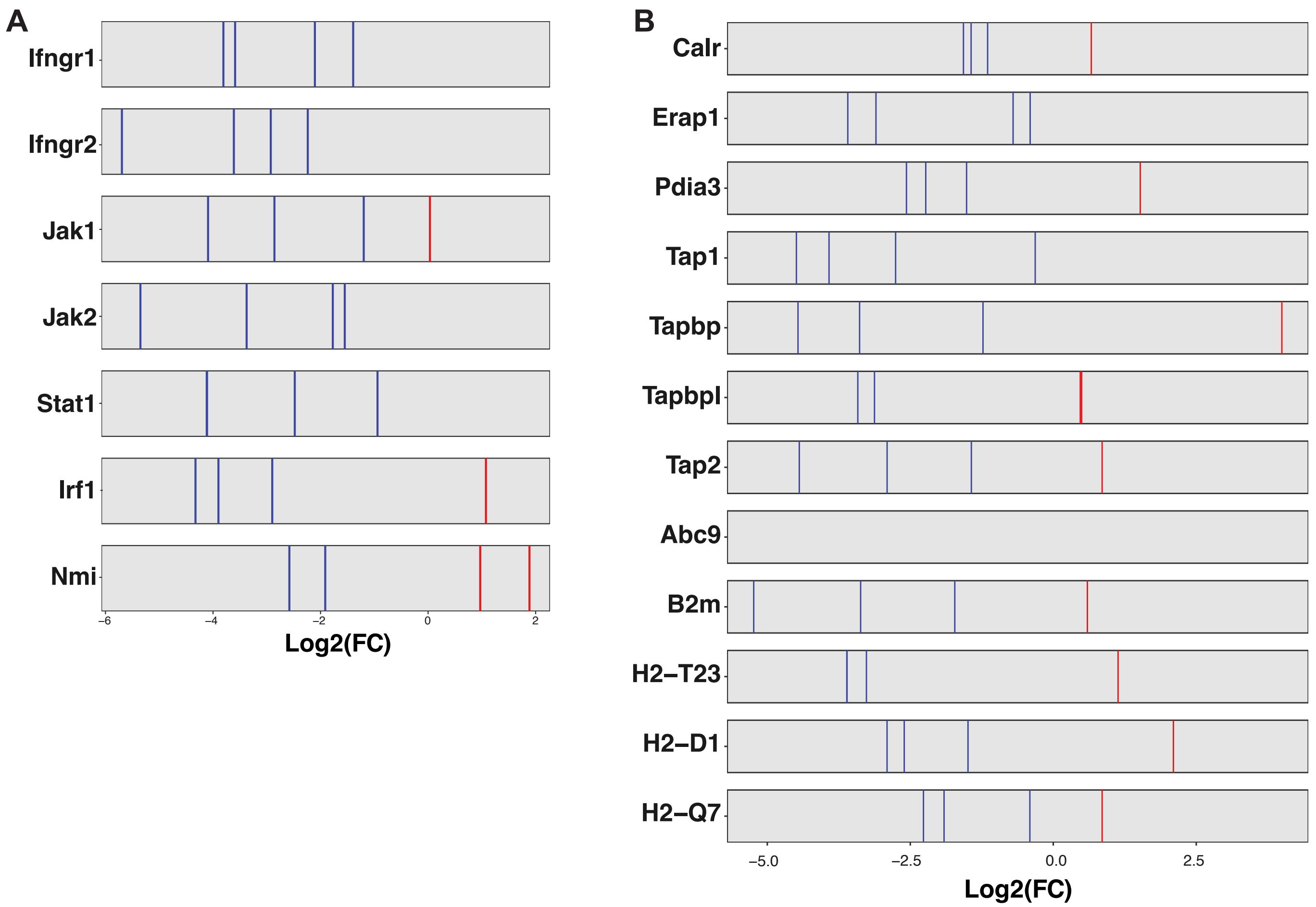

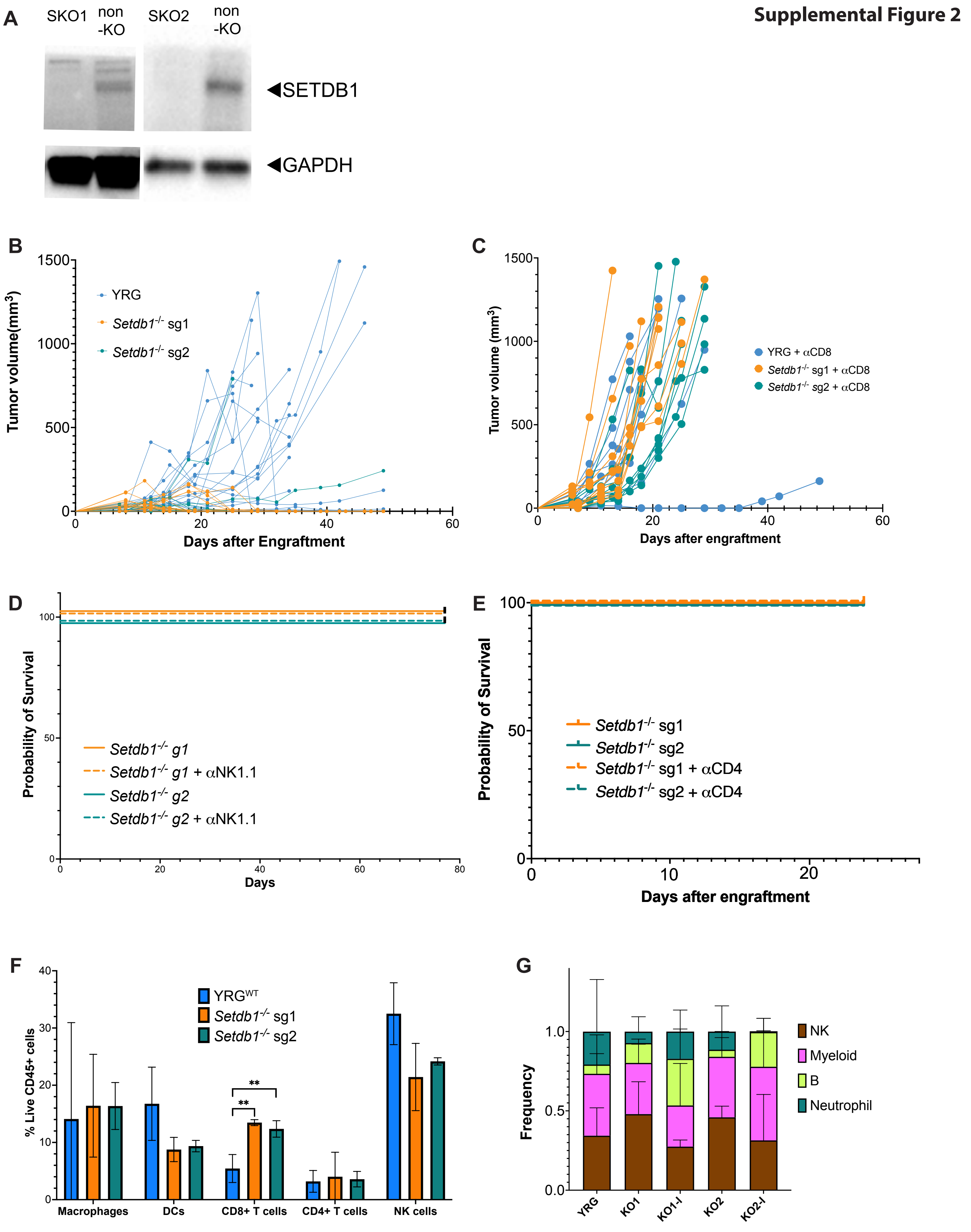

##### Supplemental Figure 3

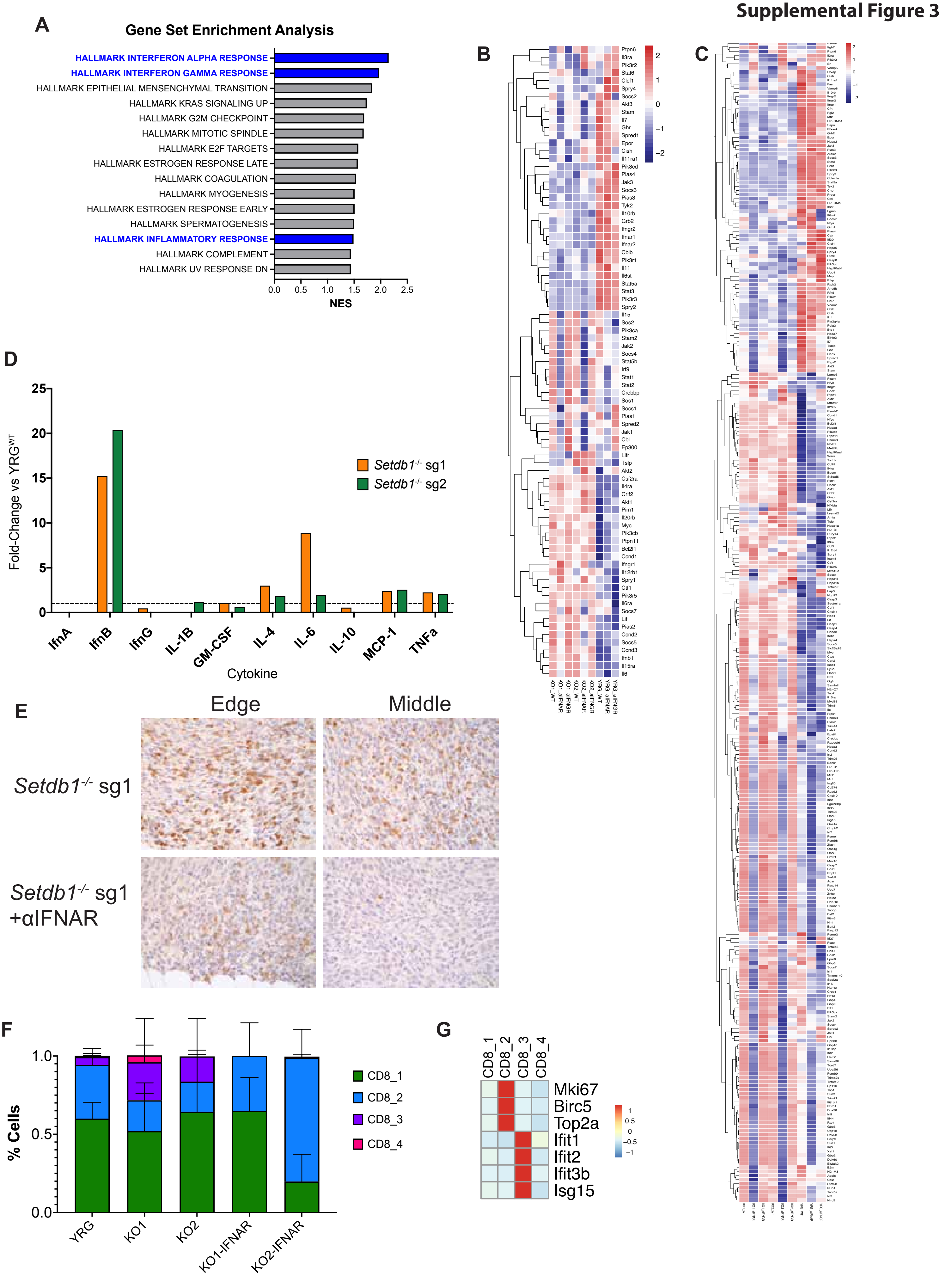

#### Supplemental Figure 4

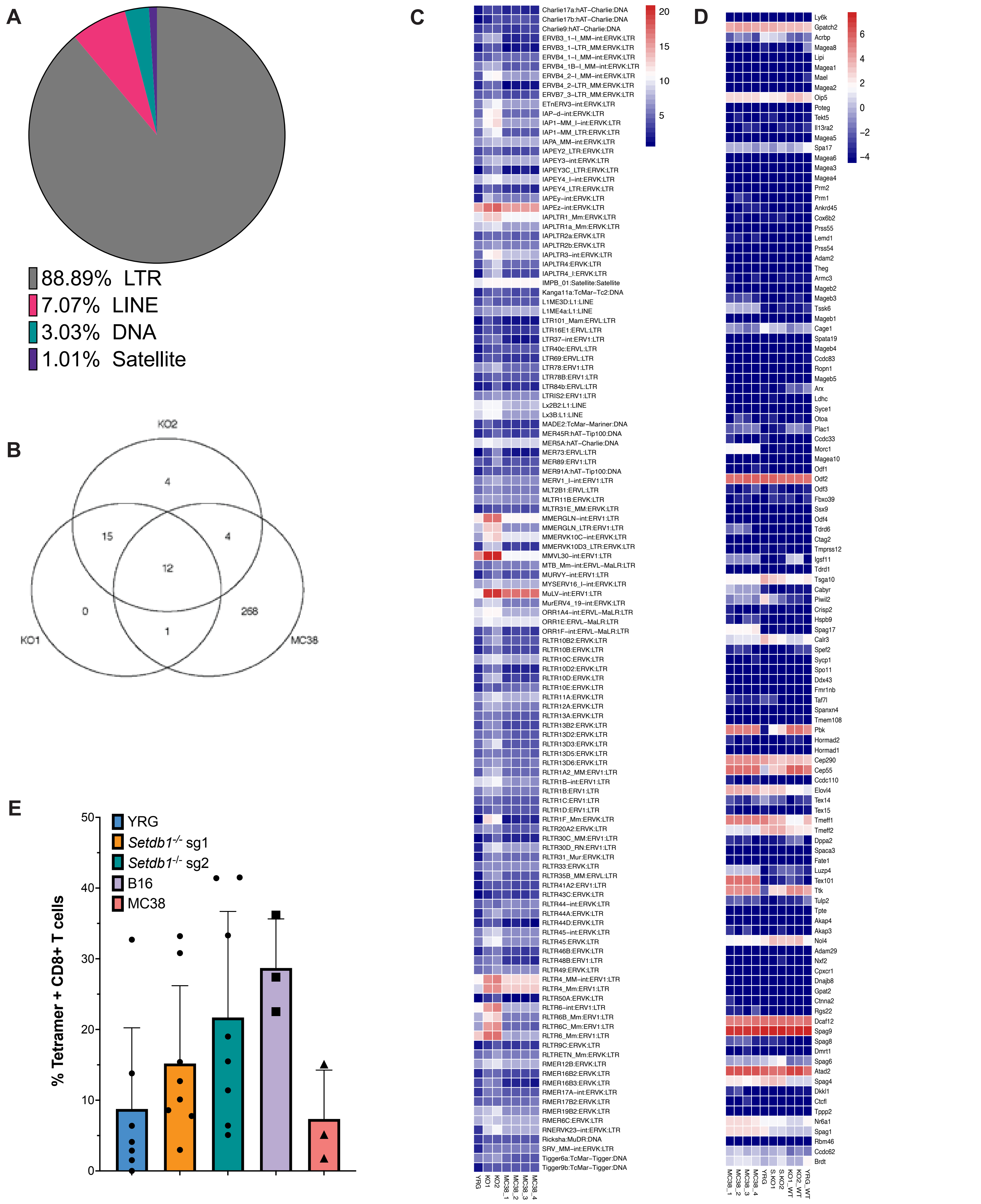

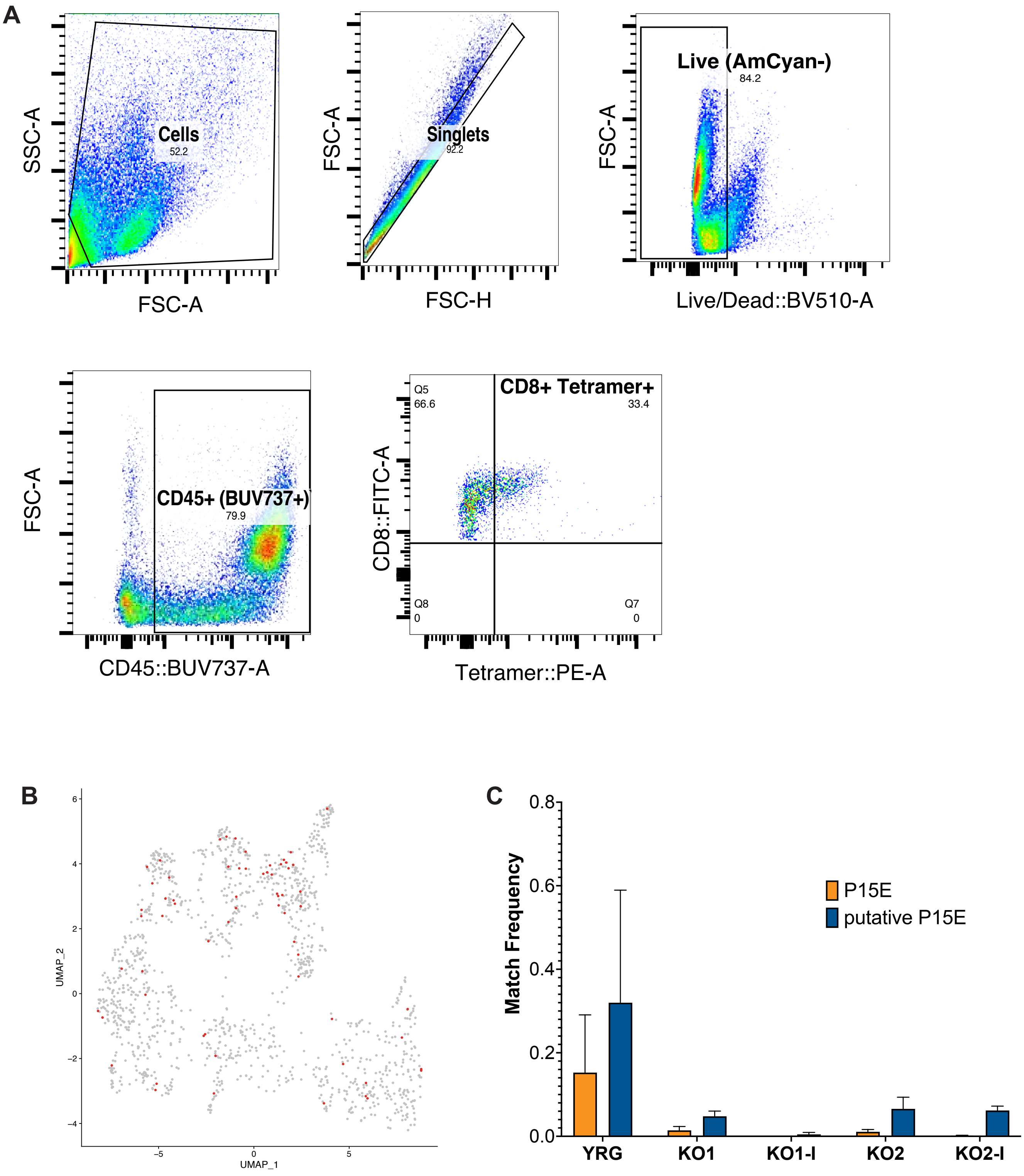
